# Physiotherapist and Podiatrist Independent Prescribing in the United Kingdom: A quasi experimental study

**DOI:** 10.1101/2020.01.09.900043

**Authors:** Nicola Carey, Judith Edwards, Simon Otter, Heather Gage, Peter Williams, Molly Courtenay, Ann Moore, Karen Stenner

## Abstract

**Background:** Increasing numbers of nurses, pharmacists and allied health professionals across the world have prescribing rights: over 90,000 of the eligible United Kingdom workforce are qualified as non-doctor prescribers. In order to inform future developments, it is important to understand the benefits and impact of prescribing by allied health professionals including physiotherapists and podiatrists.

**Aim:** to compare outcomes of Physiotherapist and Podiatrist Independent Prescriber (PP-IP) patients with those of Physiotherapist and Podiatrist non-prescribers (PP-NPs). Outcome measures included patient satisfaction, ease of access to services, quality of life and cost implications.

**Design:** a quasi-experimental, post-test control group design

**Methods:** Using mixed methods outcomes were compared between 7 sites where care was provided from a PP-IP (3 podiatrist and 4 physiotherapist IPs) and 7 sites from a PP-NP (3 podiatrist and 4 physiotherapist NPs). Patients were followed up for 2 months (2015-2016).

**Results:** 489 patients were recruited: n=243 IP sites, and n=246 NP sites. Independent prescribing was found to be highly acceptable, and equivalent in terms of quality of life (p>0.05) and patient satisfaction (p≤0.05) compared to care provided by NPs. PP-IP care delivery was found to be more resource intensive than NP-PP, with longer consultation duration for IPs (around 6.5 mins), and a higher proportion of physiotherapy patients discussed with medical colleagues (around 9.5 minutes).

**Conclusion:** This study provides new knowledge that PP-IPs provide high levels of care. PP-IP care delivery was found to be more resource intensive. Further research is required to explore cost effectiveness. A more focussed exploration within each profession using targeted outcome measures would enable a more robust comparison, inform future developments around the world and help ensure non-doctor prescribing is recognised as an effective way to alleviate shortfalls in the global workforce.

## Background

As life expectancy increases, and the world’s population continues to grow (1–3), many countries are shifting the focus of their health system from acute to chronic diseases, alongside managing increasing service demands (4). Global level predictions indicate >2 billion people will be aged >65 years by 2050, with the number > 80 years expected to double in the next decade, reaching 400 million by 2050 (1, 5). The implications for ensuring access to medicines are profound: 75% of the aging population in developed countries live with one or more chronic conditions (6), with many requiring multiple medications (5, 7). Recent data from the United Kingdom (UK), United States (US) and across Europe confirms 25% of adults take three or more medicines each day (2, 8) and that by 2020 the world’s population will receive 4.5 trillion doses of medicine each year (8–10).

There is however, a worldwide deficit of 18 million health workers (11), with a predicted 350,000 shortfall in the UK, and a third of the current workforce due to retire by 2030 (12). With a 16% increase in workload since 2010, UK workforce deficits are magnified in primary care (13), where 90% of all health encounters occur (14), and there is shortage of 2,500 general practitioners. Given the unprecedented level of future demand it is crucial that sustainable solutions that alleviate shortfalls in the global health workforce are identified (11, 12). The nature of primary care has shifted, and an increasing number of appointments in UK general practice are provided by non-medical staff (12, 15). The recent NHS Long Term Plan proposes for example, a further 20,000 non-doctor roles for primary care (16). Inadequacies with traditional doctor-led care systems mean that in order to maintain patient access to prescription medicines, new approaches are urgently required (12, 17). Allied Health Professions i.e. therapeutic radiographers, paramedics, podiatrists and physiotherapists (AHP) have in particular been identified as having an integral role to the required transformational change (18).

Extending prescribing rights to nurses, pharmacists and allied health professions (19, 20) has been the focus of a UK policy drive to improve services and access to medicines by making better use of existing skills and support service innovation (18, 21–23). Of the 907,000 UK healthcare professionals entitled to undertake prescribing training (24), over 90,000 of the eligible workforce are now qualified as prescribers (24), placing the UK as a pioneer in the development of non-doctor prescribing worldwide.

In the UK Independent Prescribing (IP) and Supplementary Prescribing (SP) are two different forms of non-doctor prescribing. Training typically involves 27 classroom days and 12 days in practice under medical supervision (25, 26), a dual qualification in IP and SP being awarded to nurses, pharmacists, radiographers and paramedics, podiatrists and physiotherapists. Independent prescribers can make prescribing decisions without the need for a doctor, while supplementary prescribing is defined as dependent prescribing, as it is based on an initial diagnosis by a doctor and an agreed clinical management plan (CMP) detailing medicines that can be prescribed by the SP (27). SP prescribing rights were extended to some allied health professions in 2005, with further changes to legislation in 2013 permitting physiotherapists and podiatrists to prescribe medicines independently (28–30).

Although several other countries, including Australia, Ireland, and Netherlands, have seen similar developments in non-medical prescribing, approaches to training, accreditation and models of prescribing practice are varied (31–34). Physiotherapists have for example, authorisation to provide advice about and/or to administer or supply medicines in some states in Australia, New Zealand and Canada, but only those in the US military can prescribe (35, 36). Podiatrists have similar authority in Australia and some European countries but are only entitled to prescribe in some Canadian states (35, 37).

When used by nurses and pharmacists, SP and IP are reported as acceptable and beneficial to patients, with some evidence of enhanced clinical outcomes compared to those achieved by doctors (32, 38–40). More recently a systematic review of non-doctor prescribing, also known as non-medical prescribing (NMP), reported that NMP has no adverse impact upon patient outcomes, patient satisfaction or resource utilisation (41). Reviews on the impact of extended physiotherapist roles reveal research hampered by small numbers of practitioners, role variation and poor role definition (42, 43), literature dominated by service descriptions and audit with positive reporting bias (35, 42, 43), and a lack of evidence regarding podiatric practice (35). Whilst PP-SP helps streamline service delivery (44, 45), IP is expected to bring additional benefits in line with nurse and pharmacist prescribing (46, 47). Exploration of clinical and cost effectiveness in this area is however limited and has to date lead to inconclusive findings (48–53). As most evidence relates to nurses and pharmacists, it is important to evaluate the impact of prescribing by allied health professionals (AHPs) in order to inform commissioning and implementation of NMP services where they are beneficial.

Six years after the introduction of current legislation enabling physiotherapists and podiatrists to prescribe independently, there has been nearly a fourfold increase in the number of physiotherapists and podiatrists with prescribing rights in England (54, 55). As of November 2019 there were 1,017 physiotherapists and 376 podiatrists with an annotation as independent prescriber, with a further 118 physiotherapists and 71 podiatrists with just supplementary prescribing (56). There is a lack of evidence of reporting on PP-IP practice, or the medicines they prescribe. Evidence from a national survey collected during preparation for the IP role indicated that PPs planned to prescribe on a regular basis, with an overall volume of prescribing suggestive of 1-2 items per day. Reflecting clinical specialities key areas of intended prescribing for physiotherapists were musculoskeletal (MSK) services, orthopaedics, respiratory and pain management, and for podiatrists’ skin, infections and MSK conditions (57).

There are additionally no studies available which quantify the impact of podiatrist and physiotherapist independent prescribing on patient satisfaction, access to services, quality of life or report cost-implications of care delivery. This is important given the increasing emphasis in the UK and around the world on extending prescribing rights to nurses, pharmacists and AHPS as a key strategy in addressing workforce deficits and ensuring patients have ongoing access to medicines (11, 12, 17, 58).

### The Study

**Aim**: was to compare the outcomes of patients managed by Physiotherapist and Podiatrist Independent Prescribers (PP-IP) with those under the care of Physiotherapist and Podiatrist non-prescribers (PP-NPs). Outcome measures included patient satisfaction, ease of access to services and quality of life. In addition, a cost-consequences analysis was undertaken which compared care delivery at the individual patient level from the NHS perspective.

### Study Design and Methods

The study adopted a quasi-experimental, post-test control group design (59). This was framed within a case study methodology used in situations when no single outcome measure is available (60, 61). Outcomes were compared between 7 sites where patients received care from a PP-IP (3 podiatrist and 4 physiotherapist IPs) and 7 sites where care was provided by a PP-NP without a prescribing qualification (3 podiatrist and 4 physiotherapist NPs) (62).

Mixed methods (including interviews, structured observation of consultations, patient questionnaires and patient record audit) were used to collect data at each of the 14 sites over 2 months (Table 1). Data collection took place simultaneously January 2015-March 2016.

**Table 1.**
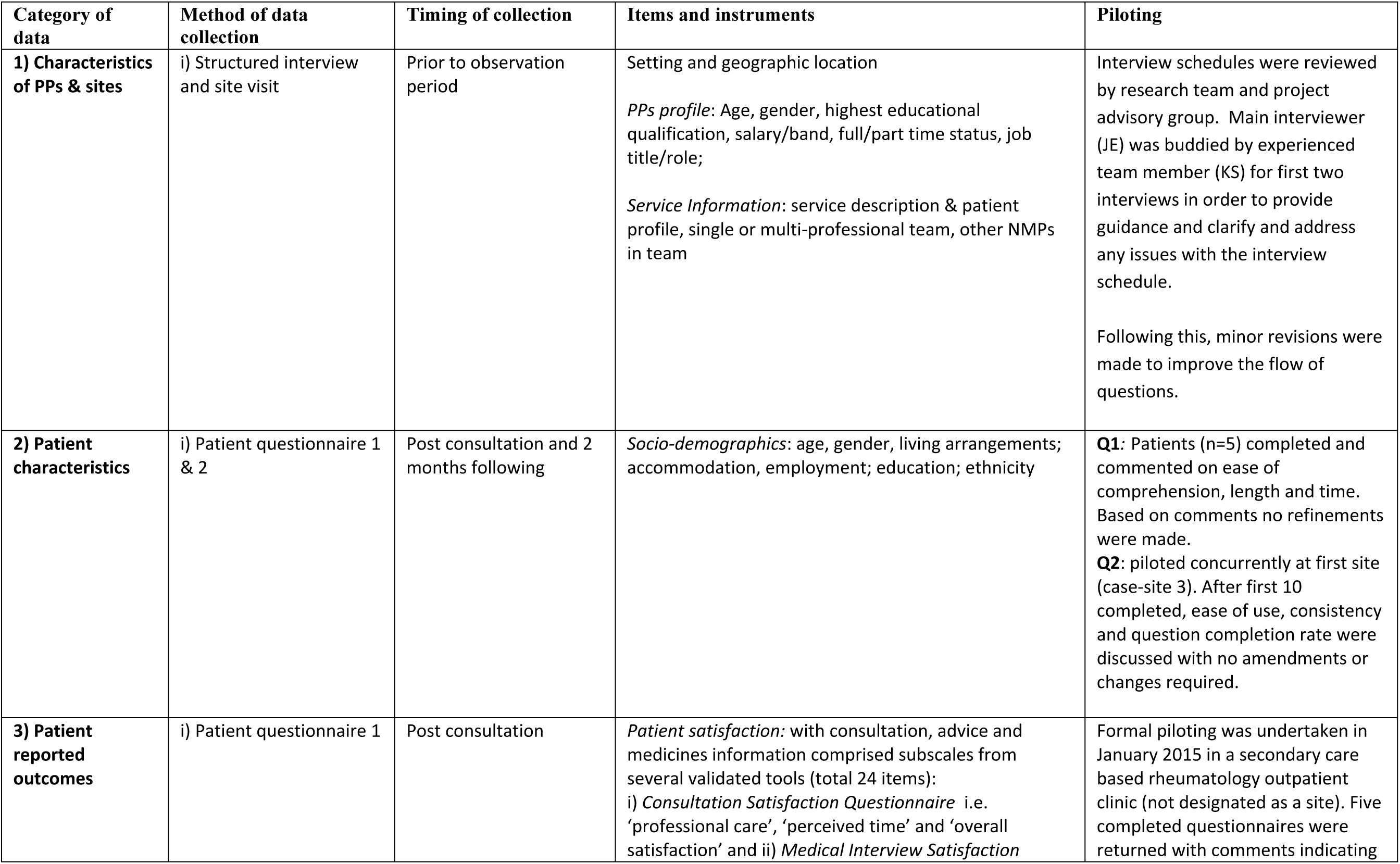
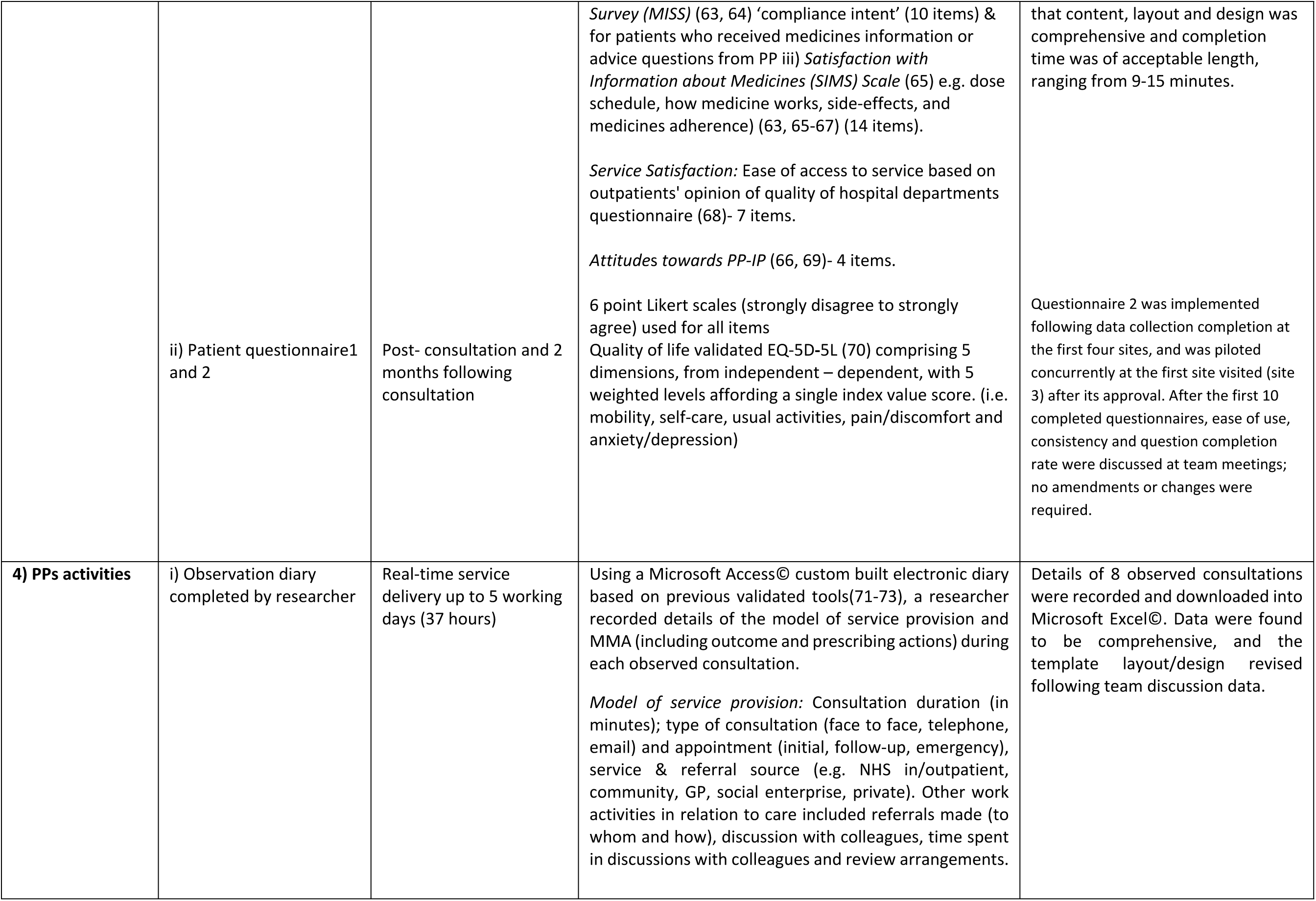
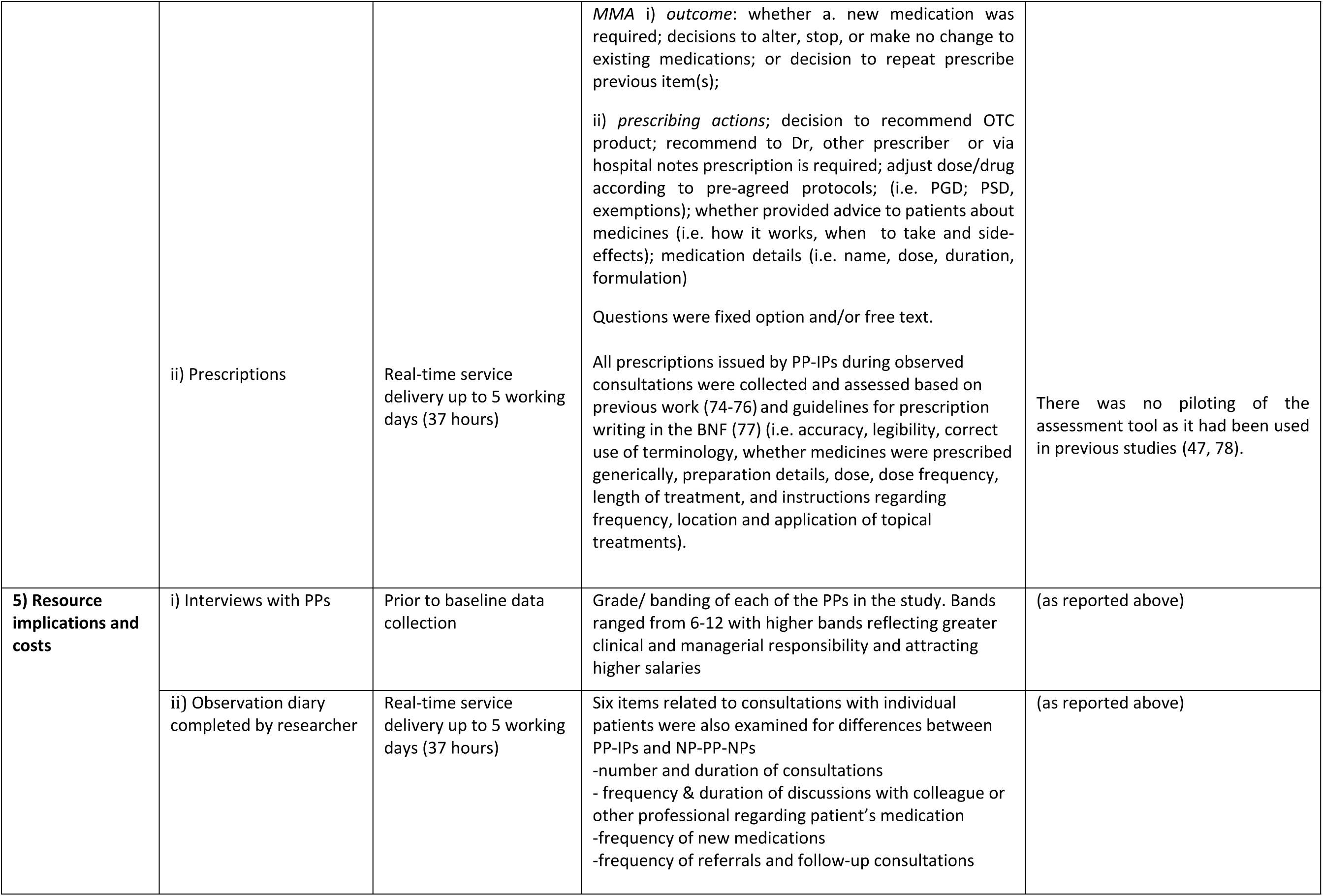
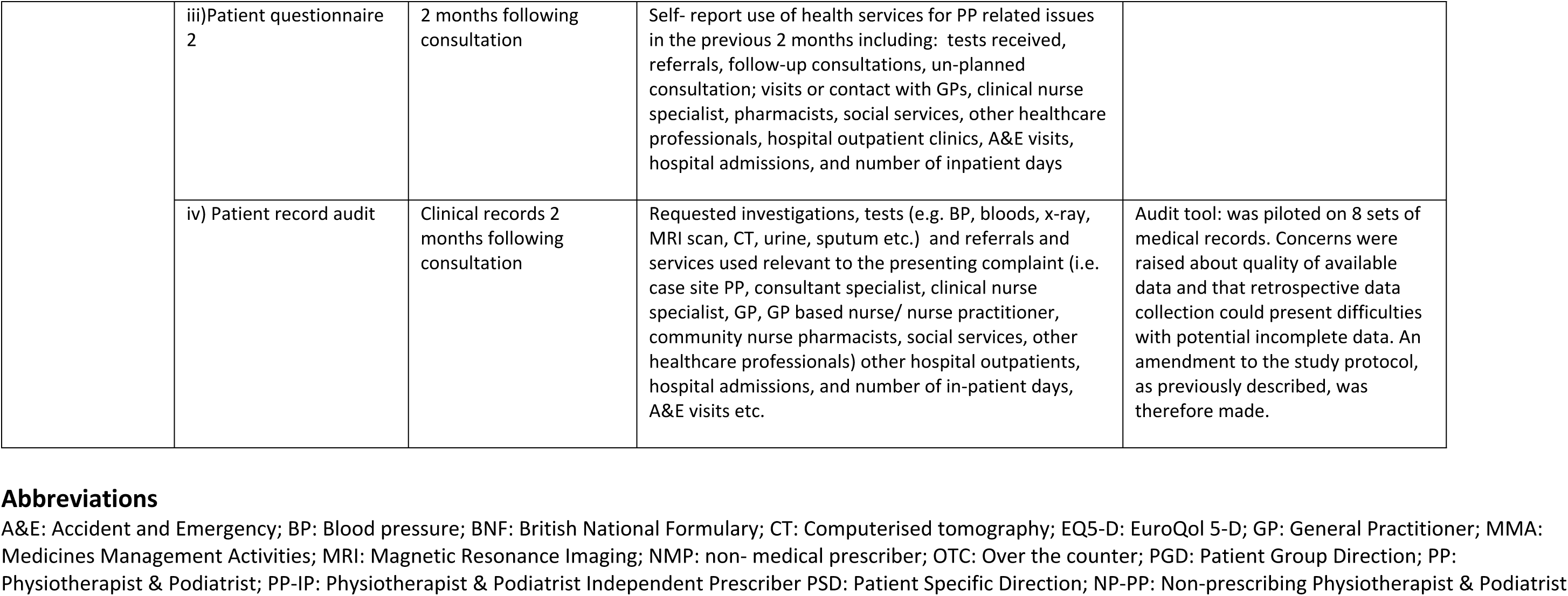
Summary of data collection arrangements and instruments.

### Sample size

The key patient outcomes related to satisfaction and ease of access to services. These were measured on a 5-point Likert scale or as Yes/No responses. The Likert scale responses were easily reducible to positive or negative responses. With respect to any dichotomous outcome variable, in order to detect an absolute underlying difference of 40% between PP-IP and NP-PPs, with size = 5% and power = 80%, a minimum of 24 subjects were needed in each PP-IP and NP-PP site. Allowing for a dropout rate of 20%, to enable a statistically sound comparison to be made between any specific pair of PP-IP and NP-PP sites, a target recruitment of 30 patients per site (total n=420), collected over a maximum of 5 working days, was set.

Initial sample estimates, based on information provided by physiotherapists and podiatrists in clinical practice, indicated that full-time PP-IPs/NP-PPs have up to 60 consultations, lasting approximately 20-40 minutes each, per week, generating data on potentially 840 patient care episodes across 14 sites, indicating that, even allowing for repeat patient visits and inclusion criteria failures, such a recruitment was feasible.

*Case sites:* Sites with PP-IPs were purposively selected from an earlier study phase (62) to include diversity with respect to care setting, geographical location and patient demographics across England.

### Recruitment

#### Podiatrists and Physiotherapists

Initial email/ telephone contact was made with PP-IPs who had completed an earlier survey whilst undertaking IP training (n=70) and indicated willingness to participate in further research (62). Those who expressed an interest were provided with a participant information sheets and supplementary information on case site involvement and requested to ensure organisational and local Research and Development (R&D) support.

Matched NP-PP sites, based on professional role, care setting, geographical location and NHS Agenda for Change (Afc) banding, were either nominated by PP-IPs, identified through personal contacts of the project advisory group or enquiries from individual Research and Development departments via the National Institute of Health Research (NIHR) portfolio. These matched NP-PPs were, with consent, contacted by a member of the research team and recruited following the same process as for PP-IPs. Written informed consent was taken from PP-IPs and NP-PPs on the first day of each case site visit by JE, who assured on-going consent with each PP-IP or NP-PP at the beginning of each contact day.

#### Patients

At each case site a consecutive sample of patients who had scheduled appointments with PP-IPs/NP-PPs during a 5-day (up to 37hrs) site visit by the study researcher (JE) were recruited in NHS sites by trained research nurses, and private sites by a second study researcher (EK) between March 2015 and February 2016.

A screening log of all patients approached for participation in the study (n=563) was recorded; both those recruited to the study (n=488, 86.7%) and those declining participation (n=75,13.3%), including hospital/unit medical record numbers, gender and the date of consent, by the local research nurse/ study researcher. Following the observed consultation (see table 1) those who agreed to participate completed and posted questionnaires into a box in the clinic area or returned using pre-paid envelopes.

### Ethical considerations

NHS Research Ethics approval from London – Surrey Borders Research Ethics Committee was (REC Ref No 14/LO/1874) and the University was obtained. R&D approval was obtained from each National Health Service (NHS) trust and private healthcare providers.

### Data collection

An initial telephone interview, informed by previous work in the area (79) was conducted with the PP from each site using semi-structured questions to gather information on site characteristics, and professional role. Details of the data collection and instruments, informed by the study patient and public involvement (PPI) and advisory groups, are presented in Table 1 All data collection instruments were piloted in a non-study physiotherapist IP NHS outpatient clinic in January 2015, with only minor corrections to wording required (see Table 1).

### Outcome measures

The patient questionnaire (Q1), informed by previous work (79) and several validated tools was designed to ensure that it: i) was relevant to patients with a range of acute and long-term conditions, attending an initial, surgical or follow-up appointment with PP: ii) supported collection of data that would allow comparisons of patient satisfaction between prescribing and non-prescribing professionals. A generic questionnaire developed to evaluate prescribing by nurses and midwives in the Republic of Ireland (66) was therefore selected for adaptation.

Two indicators of satisfaction from the post consultation questionnaire were used as outcome measures (satisfaction with the consultation and satisfaction with the advice given by the PP).

The questionnaire comprised the following subscales from validated tools:

- the subscales on ‘professional care’, ‘perceived time’ and ‘overall satisfaction’ from the Consultation Satisfaction Questionnaire (67, 80, 81)
- the ‘compliance intent’ subscale of the Medical Interview Satisfaction Survey (MISS) (63, 64)
- Questions from the Satisfaction with Information about Medicines Scale (65, 66)

**Section 1** asked participants to rate 17 statements related to patient satisfaction with services received at the time of consultation (questions 1-17). Ten questions were based on the previously validated tool Medical Interview Satisfaction scale (65, 66), and 7 additional questions designed to capture information on ‘ease of access’ to service based on outpatients’ opinion of quality of hospital departments questionnaire (68).

*Section 2a* comprised 4 statements measuring patients’ attitudes to PP-IP (66, 69) and a filter question asking whether participants had been given advice/information on medicines during the consultation. Those indicating “no” were re-directed to Section 3. Those confirming “yes” were asked to complete *section 2b*, comprising 14 statements about the advice/information they had received from PP-IPs/NP-PPs during the consultation including side effects, action of use and dose schedule and medicines adherence (63, 65–67).

*Section 3* employed the validated EQ-5D-5L quality of life profile measure of five dimensions (mobility, self-care, usual activities, pain, anxiety/ depression) rated on five levels (no problem to severe problem/ unable questionnaire (70). Although the standardized extended EQ-5D incorporates a vertical 20 cm visual analogue scale (VAS) rating scale, Patient and Public Involvement group members consistently reported difficulty indicating numerical values for how they felt at any one time point. It was therefore decided to exclude this from the questionnaire.

*Section 4* comprised 7 items related to general demographics in order to describe respondent characteristics including age, living arrangements, employment, ethnic group and educational attainment.

### Data Analysis

Quantitative data were entered in to SPSS© Version 22. Descriptive statistics were used to summarise the data and reported where open text data (specifically in relation to medication details and requested tests from the observation diary) had been converted to numeric data. When assessing change in a continuous outcome from Patient Questionnaire 1 to Questionnaire 2, a paired t-test was used.

When comparing 2 subgroups for normally distributed outcomes (notably change scores from Questionnaire 1 to Questionnaire 2, such as for overall EQ-5D-5L score), an unpaired t-test was utilised.

When comparing 2 subgroups (in particular prescribing and non-prescribing) for an ordinal outcome, a Mann-Whitney U test was utilised. When comparing 2 subgroups (notably Podiatry and Physiotherapy or prescribing and non-prescribing) for a categorical outcome, the Chi-Squared test was used, reverting to a Fisher’s Exact test in 2×2 cross tabulations if 1 or more expected cell count was found to be < 5.

#### Economic analysis

Seven resource implications of IP compared to NP were considered: rates of new prescribing; tests ordered; referrals to other health professionals; frequency of follow up; consultation duration; time spent discussing the patient with other colleagues; unplanned consultations for the same condition within two months of the index consultation. Data were gathered through the observation diary, except for tests (from the retrospective audit) and unplanned consultations (from the patient follow up questionnaire).

Group level comparisons of IP vs NP for PT and PO were undertaken separately for each of the seven variables. The cost implications (British pounds 2015) of differences in consultation length and colleague’s time spent in discussion were examined by applying nationally valid unit costs (82). A comprehensive micro level costing analysis could not be conducted because data on tests and unplanned consultations were only gathered for a sample of patients and insufficient details were available on medications, referrals and planned follow up to enable costs to be reliably ascribed. Costs that could be estimated were considered in relation to outcomes (satisfaction with consultation, satisfaction with advice, changes in health-related quality of life (EQ-5D-5L) between baseline and follow up) in a simple cost consequences framework.

## Results

### Characteristics of participants

#### i) PPs and case sites

Seven matched pairs of sites, (3 podiatry and 4 physiotherapy) were recruited. Sites were based across 8 Academic Health Science Networks in England (https://www.ahsnnetwork.com/), a mixed range of settings, including private practice (n=2), primary care (n=6), secondary care (n=6), social enterprise (n=2) and were well matched by professional role, care setting and agenda for change banding (see Table 2). All PP-IPs had been qualified for at least 12 months prior to data collection. A total of 489 patients were recruited: 243 across the PP-IP sites with 246 across the NP-PP sites.

**Table 2:** Characteristics of the sites and Physiotherapists and Podiatrists.

| Pair | Case study site | No. Patients Recruited | Type of PP | Job Title | Setting | Location in England * | Age | Salary band | Full or part time <30 hrs in practice | Education highest | Single or multi-professional team | Patient questionnaire 1 | Follow up-Patient Questionnaire | Prescriptions |
| --- | --- | --- | --- | --- | --- | --- | --- | --- | --- | --- | --- | --- | --- | --- |
| 1 | 1 | 49 | PO-IP | General/Private | Private | London | 71 | 8a | Full time | Doctorate | single | 40 | N/A | 0 |
|  | 2 | 46 | PO-NP | General/Private | Private | London | 47 | 12 | Full time | Masters | single | 35 | N/A | n/a |
| 2 | 3 | 33 | PO-IP | Specialist | Secondary care, NHS In/out patient | Wessex | 41 | 7 | Full time | Masters | multi-professional | 22 | 19 | 6 |
|  | 8 | 37 | PO-NP | Specialist | NHS primary & secondary (& private) | Kent, Surrey, Sussex | 39 | 6 | Full-time | Degree | single | 25 | 22 | n/a |
| 3 | 10 | 51 | PO-IP | Surgeon/consultant | NHS secondary (& private) | Oxford | 59 | 9 | Full time | Masters | multi-professional | 32 | 38 | 3 |
|  | 6 | 42 | PO-NP | Surgeon/consultant | NHS secondary | North East & North Cumbria | 47 | 9 | Part-time | Masters | multi-professional | 26 | 23 | n/a |
| 4 | 7 | 6 | PT-IP | Specialist | Community | London | 31 | 7 | Part-time | Masters | multi-professional | 25 | N/A | 0 |
|  | 4 | 11 | PT-NP | Specialist | NHS Primary, Community care | Kent, Surrey, Sussex | 47 | 8a | Full time | Masters | multi-professional | 25 | N/A | n/a |
| 5 | 9 | 42 | PT-IP | Specialist | Primary, community Social enterprise | Kent, Surrey, Sussex | 46 | 8a | Full time | Diploma | multi-professional | 2 | 2 | 3 |
|  | 5 | 38 | PT-NP | Surgeon/consultant | Tier 2 NHS ESP assessment service | Wessex | 42 | 8a | Part-time | Doctorate | multi-professional | 6 | 3 | n/a |
| 6 | 11 | 41 | PT-IP | Specialist | Acute Foundation Trust | Northwest coast | 58 | 8a | Full time | Masters | multi-professional | 27 | 29 | 0 |
|  | 12 | 35 | PT-NP | Surgeon/consultant | NHS secondary care | Kent, Surrey, Sussex | 48 | 8c | Full time | Masters | multi-professional | 19 | 23 | n/a |
| 7 | 13 | 21 | PT-IP | Specialist | NHS primary & community Social enterprise | Kent, Surrey, Sussex | 52 | 8a | Full time | Masters | multi-professional | 8 | 16 | 3 |
|  | 14 | 36 | PT-NP | Specialist | Primary & community | Kent, Surrey, Sussex | 38 | 8a | Full time | Masters | multi-professional | 23 | 20 | n/a |
|  |  |  |  |  | Social enterprise |  |  |  |  |  |  |  |  |  |
| **Totals** |  | **488** |  |  |  |  |  |  |  |  |  | **315** | **195** | **15** |

PO-IP: Podiatrist independent prescriber PO-NP: Podiatrist non-prescriber PT-IP: Physiotherapist independent prescriber PT-NP: Physiotherapist non-prescriber
Job title: **Surgeon/Consultant** (consultant physiotherapists, consultant podiatric surgeons) **General/Private** (physiotherapists practitioners, physiotherapists, podiatrists) **Specialist** (e.g. Clinical specialist physiotherapists, extended scope physiotherapist clinical specialist podiatrists, clinical lead or senior podiatrists)

Nearly all consultations (n=474), both PP-IPs and PP-NPS, were face to face (n=473, 99.8%), duration 2-203 minutes. There was considerable variation in the location of services: 39.2% (n=186) of consultations were provided in NHS hospital outpatients, 25.1% (n=119) NHS community clinics, 20.3% (n=96) private practice, 9.7% (n=46) general practice, 4.4% (n=21) social enterprise and 1.3% (n=6) community service.

#### ii) Patients

Demographic data (see table 3) were collected from 315/ 468 (67.3%) patients who consented to and returned the initial questionnaire: 49.5% (n=156) were from prescribing and 50.5% (n=159) from non-prescribing sites. The samples were similar in terms of age, employment status, level of formal education, and ethnic group (p>0.05).

**Table 3:**
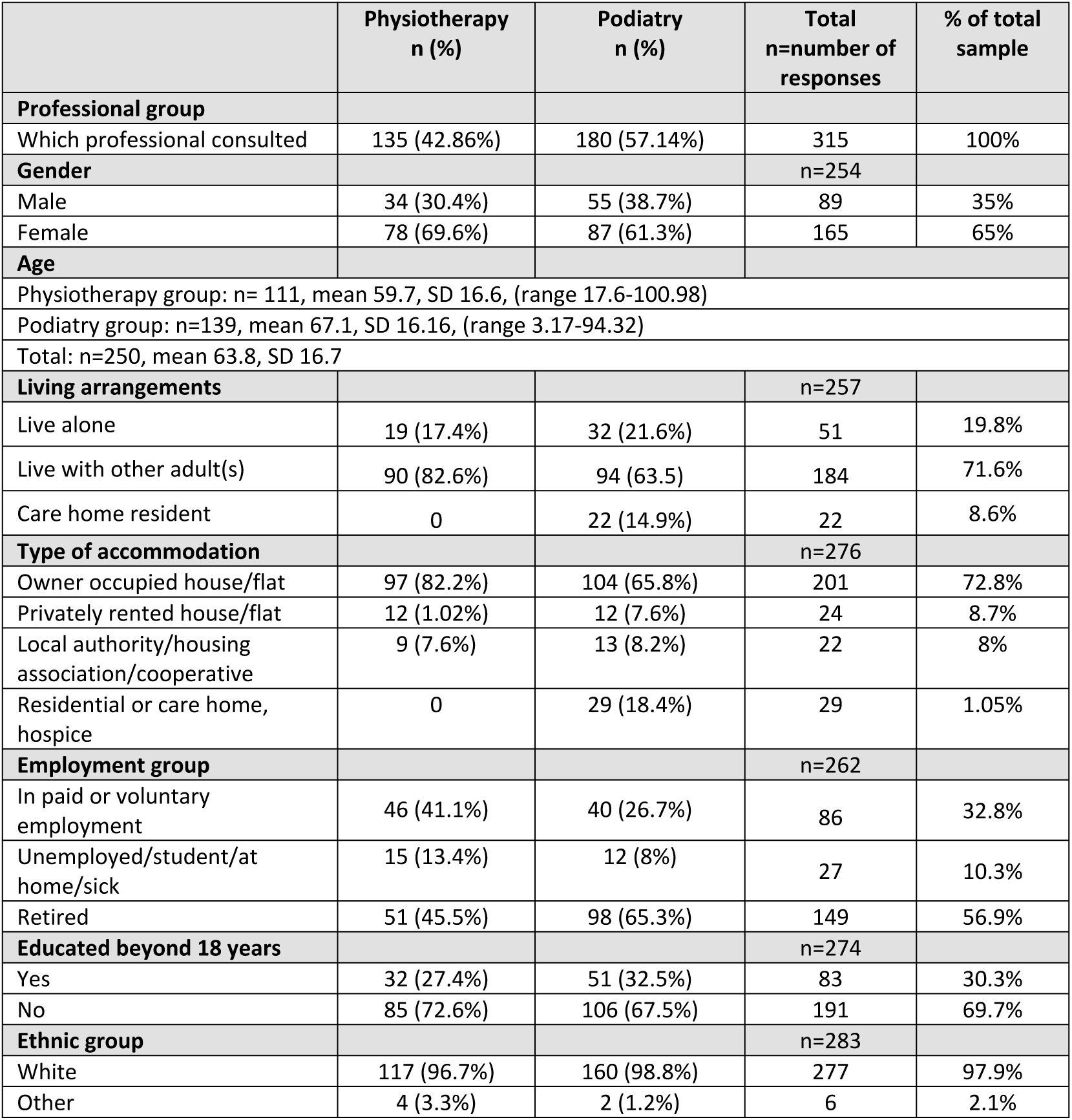
Patient characteristics.

#### iii) Patient outcomes

##### a) Satisfaction and access to services

The majority of patients (75.9%, n=239) agreed that PP’s should be able to prescribe medicines for patients, however 23.2% (n=73) would prefer a doctor to prescribe. Levels of satisfaction for the sample as a total were high, with over 60% positive agreement on all items other than ability to contact the service in an emergency (n=144, 44.4%). Satisfaction with 17 specified aspects of the consultation and services provided by PPs indicated a significantly higher level of satisfaction among the patients of PP-IPs than those of NP-PPs in 8 instances (table 4).

**Table 4.**
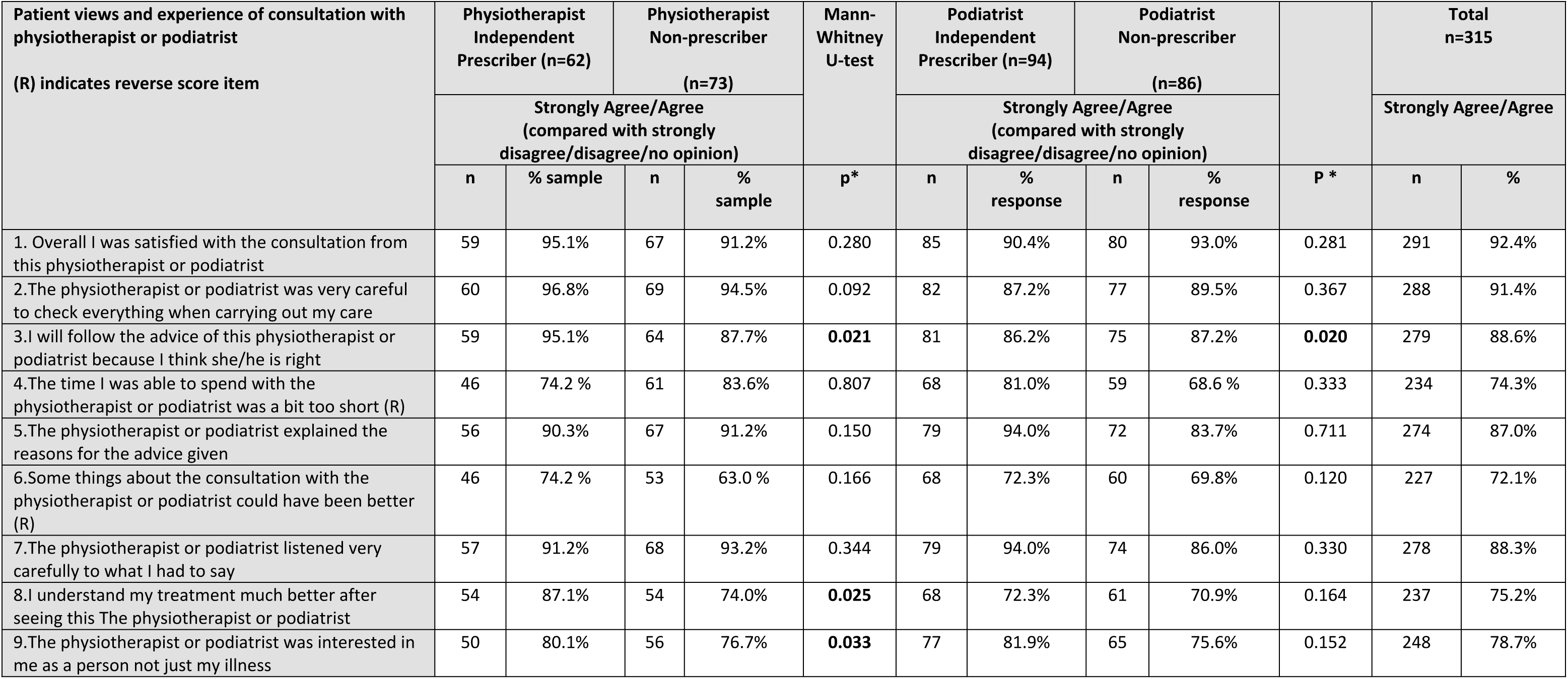
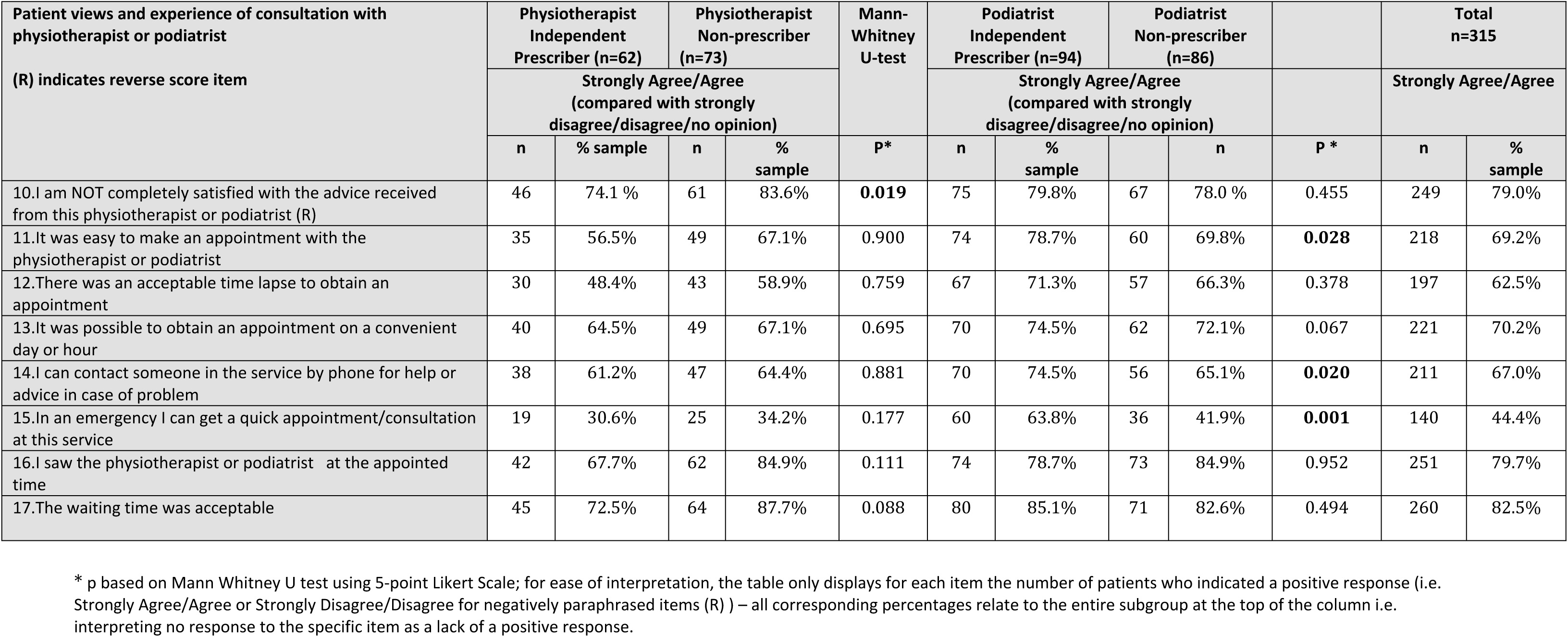
Patient views and experience of satisfaction with care received from physiotherapist or podiatrist.

With respect to service access, patients of PO-IPs were more satisfied with ‘the ease of making an appointment’ and ‘the ability to contact the service by phone or in times of emergency’ (see table 4) than NP-PO patients, with no notable difference evident in patients attending PT-IPs compared to NP-PTs.

There was no effect on the remaining four items reporting on ease of access on the acceptability of: i) waiting time to obtain an appointment; ii) obtaining an appointment on a convenient day or hour; iii) waiting time or iv) seeing the PT or PO at the appointed time between patients attending a PP-IPs when compared to those attending a NP-PP.

Patients of PP-IPs were more likely to receive medicines information or advice during the consultation (58 out of 146 (39.7%) vs 37 out of 151 PP-NP patients (24.5%); p=0.005), with varying levels of satisfaction reported (see table 5). Compared to PT-NPs patients, PT-IP patients were significantly more likely to: ‘be told when’ and ‘how often’ to take their medicine, ‘intend to take their medicines’ and ‘find it easier to follow the PT advice’ (p≤ 0.05).

**Table 5.**
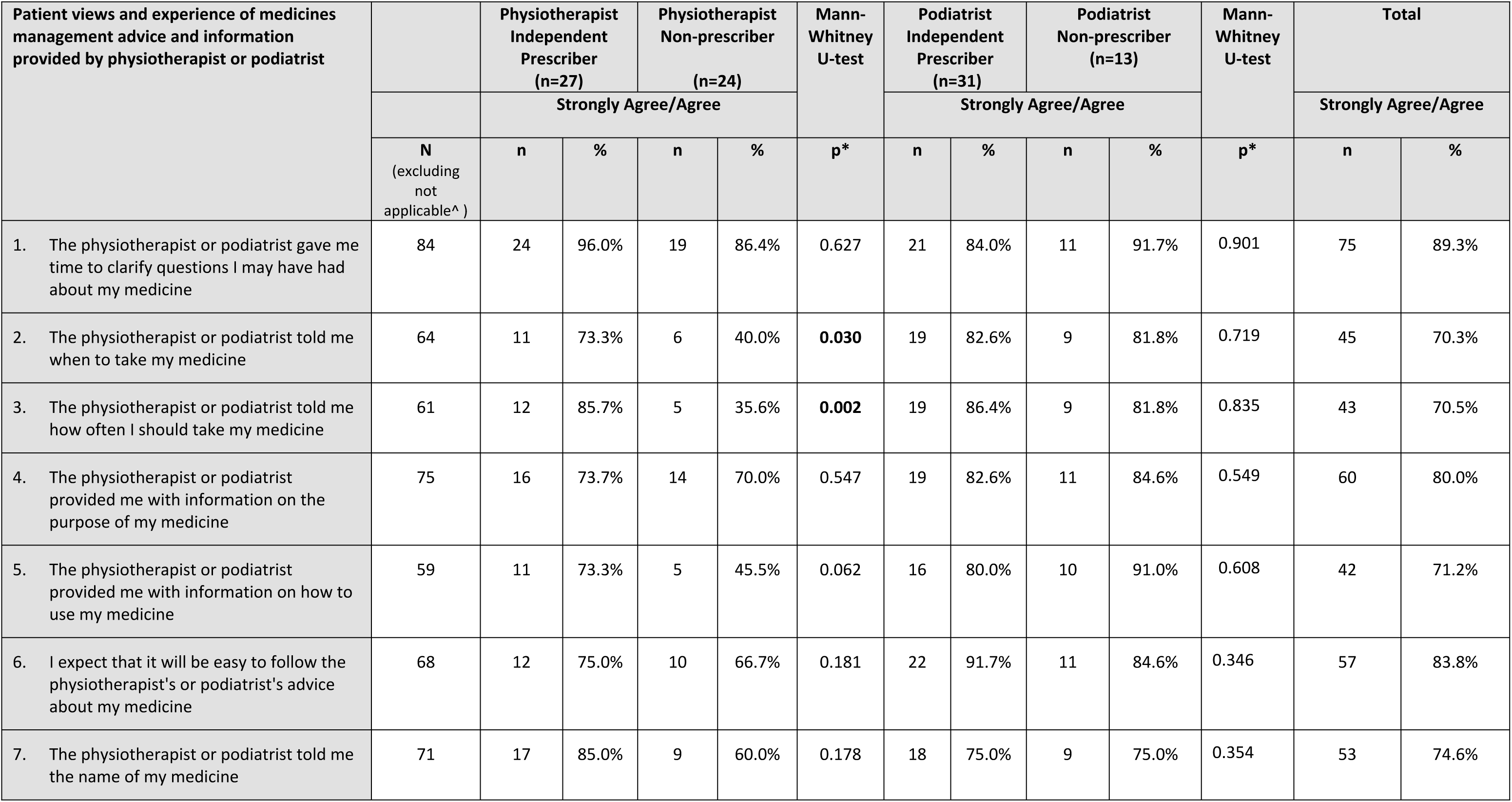
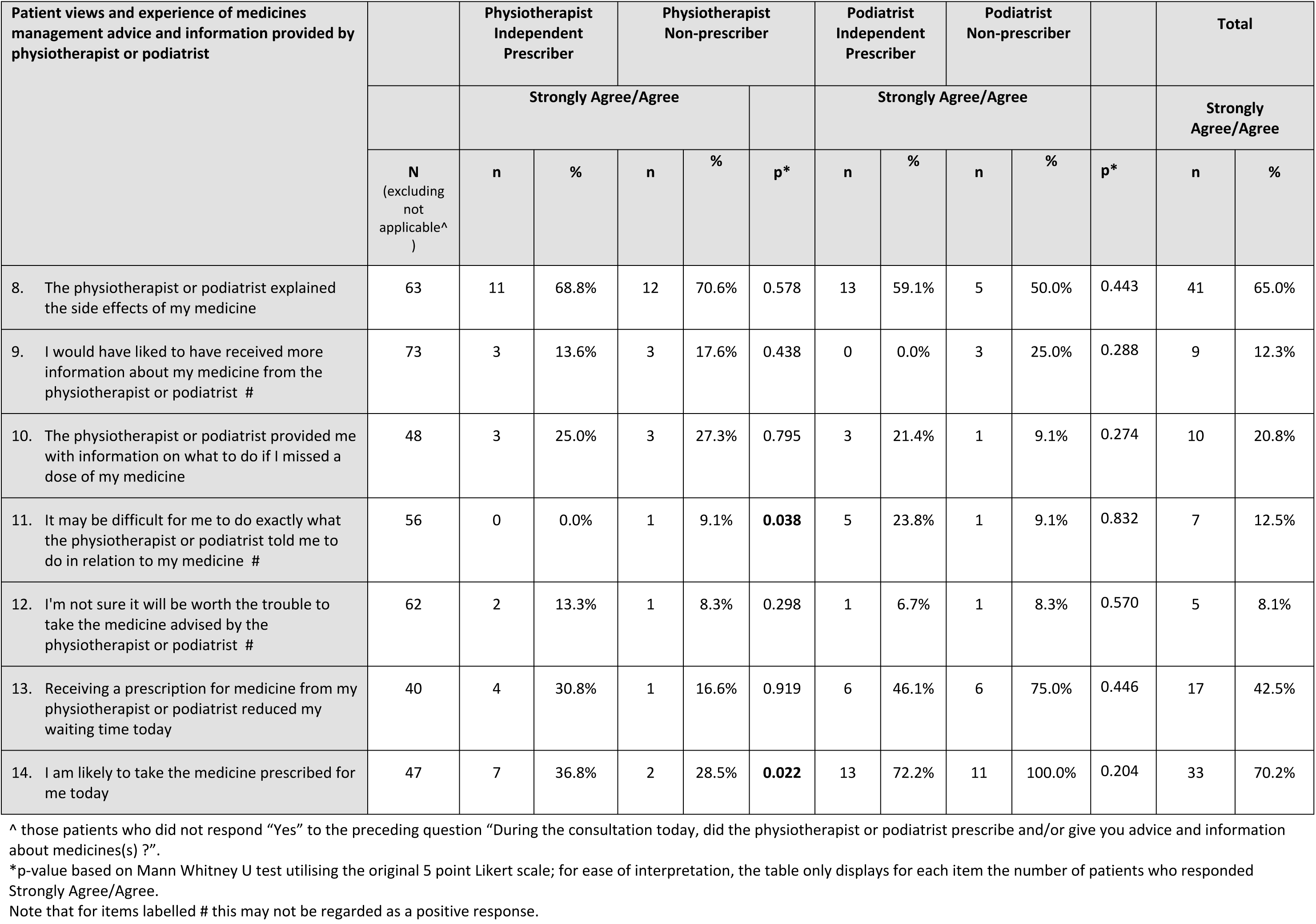
Patient views and experience of medicines management advice and information provided by physiotherapist or podiatrist.

##### b) Quality of life-EQ-5D-L

Indications at baseline were that patients who saw PP-IPs scored less for mobility, however there was no statistically significant difference between PP-IP and PP-NP groups on either individual items or overall score (p≥0.05).

Indications at baseline were that patients who saw PT-IP had lower generic quality of life than those seeing the PT-NP, due to lower scores on the mobility dimension. However, there was no statistically significant difference between PP-IP and PP-NP groups on either individual items or overall EQ-5D-5Lscore (p≥0.05) (Table 6, individual dimension scores not shown).

**Table 6:** Overall EQ5D index score: baseline and follow-up.

|  | From the 129 completers | Baseline for 116 with EQ5D in BOTH data sets only | Follow-Up for 116 with EQ5D in BOTH data sets only |  |  |
| --- | --- | --- | --- | --- | --- |
|  | Number of patients completing BOTH sets of EQ5D questions | EQ5D-5L Mean (SD) | EQ5D-5L Mean (SD) | Change from Baseline (95% CI)* | Paired t-test p-value |
| PT IP | 25 | 0.56 (0.31) | 0.64 (0.27) | 0.08 (-0.04 to 0.19) | 0.194 |
| PT NP | 28 | 0.73 (0.19) | 0.73 (0.22) | 0.001 (-0.07 to 0.07) | 0.973 |
| PO IP | 33 | 0.70 (0.26) | 0.78 (0.20) | 0.08 (0.003 to 0.16) | 0.042 |
| PO NP | 30 | 0.66 (0.26) | 0.76 (0.28) | 0.10 (0.03 to 0.16) | 0.004 |
| All IP | 58 | 0.64 (0.29) | 0.72 (0.24) | 0.08 (0.01 to 0.14) | 0.019 |
| All NP | 58 | 0.69 (0.23) | 0.75 (0.25) | 0.05 (0.003 to 0.10) | 0.036 |
| All PT | 53 | 0.65 (0.26) | 0.69 (0.25) | 0.04 (-0.03 to 0.10) | 0.266 |
| All PO | 63 | 0.68 (0.26) | 0.77 (0.24) | 0.09 (0.04 to 0.14) | 0.001 |

\*[Positive change indicates mean improvement in health at Follow-Up]

Quality of life overall scores in both PP-IP and PP-NP groups improved significantly between baseline and follow-up. Differences in change scores between the PP-IP and PP-NP group, however, were not statistically significant (Table 6). The sample for which data at both time points were available was limited (n=116).

#### Economic analysis-patient level care delivery

*Physiotherapists*: Compared to NPs, the physiotherapist IPs had significantly longer consultation duration (20.8 vs 27.6 minutes) (Table 7).

**Table 7.** Comparison of independent prescribers and non-prescribers, by profession, on variables used in the cost analysis.

| Professional group | Prescribing status |  | Number of new medications required (Observation Q6) | Number of tests requested / patient (Sample audit) | Consultation time in minutes / patient (Observation Q1) | Discussions with colleagues in minutes/ per patient (Observation Q9,10) | x | Patients receiving referral (not for tests) (Observation Q11) | Patients with planned follow up (Observation Q15) | Patients reporting verified unplanned consultations within 2 months (Patient questionnaire) |
| --- | --- | --- | --- | --- | --- | --- | --- | --- | --- | --- |
| PHYSIO-THERAPY | Independent prescriber (IP) | N | 107 | 42 | 107 | 107 | N | 107 | 107 | 47 |
|  |  | Missing | 9 | 74 | 9 | 9 | Yes N | 32 | 54 (8 by phone) | 1 |
|  |  | N, % of zeros | 75, 70.1% | 32, 76.2% | 0 | 88, 82.2% | Yes % | 29.9% | 50.5% | 2.1% |
|  |  | Mean | 0.327 | 0.262 | 27.64 | 1.802 |  |  |  |  |
|  |  | SD | 0.546 | 0.497 | 14.10 | 5.585 |  |  |  |  |
|  |  | Median | 0 | 0 | 24 | 0 |  |  |  |  |
|  |  | IQR | 0 to 1 | 0 to 0.25 | 18 to 34 | 0 to 0 |  |  |  |  |
|  | Non prescriber (NP) | N | 115 | 44 | 115 | 115 | N | 115 | 115 | 46 |
|  |  | Missing | 7 | 78 | 7 | 7 | Yes N | 34 | 51 (1 by phone) | 2 |
|  |  | N, % of zeros | 87, 75.7% | 33, 75.0% | 0 | 114, 99.1% | Yes % | 29.6% | 44.3% | 4.3% |
|  |  | Mean | 0.252 | 0.250 | 20.83 | 0~ |  |  |  |  |
|  |  | SD | 0.456 | 0.438 | 10.46 | 0~ |  |  |  |  |
|  |  | Median | 0 | 0 | 19 | 0 |  |  |  |  |
|  |  | IQR | 0 to 0 | 0 to 0.75 | 14 to 28 | 0 to 0 |  |  |  |  |
|  | Significant difference (p) |  | MWU 0.336 | MWU 0.949 | MWU <0.0005 | MWU <0.0005 |  | Chi Sq 0.956 | Chi Sq 0.361 | FE 0.617 |
| PODIATRY | Independent prescriber (IP) | N | 128 | 24 | 128 | 128 | N | 128 | 128 | 57 |
|  |  | Missing | 5 | 109 | 5 | 5 | Yes N | 17 | 110 (0 by phone) | 0 |
|  |  | N, % of zeros | 93, 72.7% | 17, 70.8% | 0 | 109, 85.2% | Yes % | 13.3% | 85.9% | 0% |
|  |  | Mean | 0.328 | 0.375 | 24.27 | 0.976 |  |  |  |  |
|  |  | SD | 0.616 | 0.647 | 24.32 | 2.682 |  |  |  |  |
|  |  | Median | 0 | 0 | 16 | 0 |  |  |  |  |
|  |  | IQR | 0 to 1 | 0 to 1 | 11 to 27.75 | 0 to 0 |  |  |  |  |
|  | Non prescriber (NP) | N | 124 | 32 | 123 | 124 | N | 124 | 124 | 47 |
|  |  | Missing | 3 | 95 | 4 | 3 | Yes N | 6 | 111 (7 by phone) | 1 |
|  |  | N, % of zeros | 114, 91.9% | 32, 100% | 0 | 111, 89.5% | Yes % | 4.8% | 89.5% | 2.1% |
|  |  | Mean | 0.105 | 0 | 16.88 | 0.726 |  |  |  |  |
|  |  | SD | 0.379 | 0 | 9.86 | 2.867 |  |  |  |  |
|  |  | Median | 0 | 0 | 16 | 0 |  |  |  |  |
|  |  | IQR | 0 to 0 | 0 to 0 | 10 to 23 | 0 to 0 |  |  |  |  |
|  | Significant difference (p) |  | MWU 0.001 | MWU <0.0005 | MWU 0.073 | MWU 0.349 |  | Chi Sq 0.20 | Chi Sq 0.387 | FE 0.452 |

*Podiatrists:* The frequency with which new medications and tests were ordered were significantly higher in IP than NP (Table 7). There was a trend for consultation duration to be longer for IP (23.4 vs 19.9 minutes) (Table 7).

Planning of follow up consultations was higher by podiatrist IPs than physiotherapist IPs, but no significant differences were found between IP and NP within the professions. After removing unplanned consultations in the two months after the original consultation that were considered (by two independent reviewers) to be unamenable to treatment delivered in the index consultation, only four items of unplanned service utilisation remained across the whole sample of patients of PT and PO, all of which were related to pain relief (Table 7).

#### Costs

Difference in costs of consultation duration of IP v NP for physiotherapist and podiatrist groups were based on Agenda for Change (AfC) band 8a, which was the most frequent grade of PP-IPs in the study, i.e. £70 per hour (82). Compared to the cost of a NP consultation, the IP consultation was, on average, more costly by £7.95 for PT (£24.30 vs £32.25) and £8.62 (£19.69 vs £28.31) for PO. The salary of a grade 9 professional is twice that of grade 8a, so at that higher level, the differences in the cost of consultations between IP and NP would be doubled. Use of grade 7 instead of grade 8a would reduce the differences between IP and NP by about £1.20 per consultation. Amongst the POs, the IPs were at band 7 (advanced / team leader), 8a (principal) and 9 (consultant); two of the NPs were band 9 and the third was band 6 (specialist). Participating physiotherapists were all band 8a, except one NP (grade 8c), and one IP (grade 7)

Costs could not be estimated for the other elements of activity that might differ between IP and NP due to data problems. Information on tests ordered were drawn from a small sample of records in each site (the audit); reporting of the type and dose of new medications, referrals and frequency of planned follow up was incomplete.

#### Discussions with colleagues

The IPs in the PT group consulted colleagues about patients significantly more often than the NPs (17.8% vs 0.9% of consultations), and most discussions were with medical colleagues, averaging 9.5 minutes per discussion (Table 8).

**Table 8.** Discussion with colleagues about patient.

| Professional group | Prescribing status | Number and % of all patients seen for whom discussion occurred with colleague | Mean (SD) minutes in discussions with colleague per patient | Discussion with same professional n, mean (SD) minutes | Same colleague cost / discussion* (£, 2015) | Discussion with medical professional n, mean(SD) minutes | Medical colleague cost / discussion* (£, 2015) |
| --- | --- | --- | --- | --- | --- | --- | --- |
| PHYSIOTHERAPY | Independent prescriber | 19 (17.8%) | 10.61 (9.68) | 3, 19.5 (14.8) | £22.75 | 16, 9.5 (8.9) | £21.69 |
|  | Non prescriber | 1 (0.9%) | 0 (n/a) | 1, time missing | Not known | 0, n/a | 0 |
|  | Significant difference | p<0.0005# | n/a |  |  |  |  |
| PODIATRY | Independent prescriber | 19 (14.8%) | 6.89 (3.20) | 11, 6.8 (3.6) | £7.93 | 8, 7.0 (2.8) | £15.98 |
|  | Non prescriber | 13 (10.5%) | 6.92 (6.14) | 12, 7.3 (6.3) | £8.52 | 1, 3.0 (0.0) | £6.85 |
|  | Significant difference | p=0.299~ | p=0.493^ |  |  |  |  |
| # Fishers Exact test; ~ Chi squared test; ^ Mann Whitney U test |  |  |  |  |  |  |  |
| * Unit costs of health and social care 2015 (Curtis and Burns 2015), pro rata based on £70/ hour for same professional i.e. AfC band 8a, as in Ec2 above, and £137/ hour for medical consultant |  |  |  |  |  |  |  |

POs held discussions with colleagues for >10% of consultations (14.8% IPs, 10.5% NPs, (Table 8)), for around 7 minutes. IPs discussed a higher proportion of patients with medical colleagues, than a colleague from the same profession, thereby likely to be incurring higher costs. However, information on colleagues consulted was not precise, so calculations were indicative only. Some POs were band 9 (consultant), so reporting discussions with ‘same’ professional would imply higher costs than are indicated in the table, which are based on AfC band 8a.

#### Cost consequences analysis

The available data suggest that for both PT and PO in this study, care delivery by IP is more resource intensive and costly than NP due to longer consultations for PTs, and taking more time of colleagues to discuss patients. Whilst not costed, PO-IPs had higher frequency of ordering medications and tests than PO-NPs. Analysis of the changes in self-reported health status between baseline and 2 month follow up using EQ-5D-5L found no difference in change scores of IP and NP for either PT or PO, but these data were only available for a small sample of participants.

## Discussion

This is the only known national evaluation of PP-IP in the UK or the world, and the first to adopt a quasi-experimental research design to compare outcomes and costs for patients managed by PP-IPs and NP-PPs. Unlike nurses and pharmacists, where prescribing has been explored in some detail using self-reported outcomes (32, 53, 79), there is a dearth of equivalent information in the allied health professions, including either physiotherapy and/ or podiatry (35, 41) and/ or studies adopting direct observation of outcomes (32). Our study demonstrates that care provided by PP-IPs is equivalent, in terms of quality of life and patient satisfaction, to care provided by NP-PPs with prescribing undertaken by doctors. IP by PPs was found to be highly acceptable, with higher levels of patient satisfaction in some aspects of medicines information also reported than for NPs.

Importantly, it appears that PP-IP is developing in line with original policy intention to improve access and quality of care in across a range of settings (83–85). The evidence generated in this study demonstrates that PP-IPs can provide a high standard of care. Extending non-medical staff, such as physiotherapists’ and podiatrists’, scope of practice to include independent prescribing is key to supporting effective delivering of the NHS Long Term Plan (12, 58, 86), and in creating a step change in developing the capacity and capability of the workforce to deliver innovative models of service delivery (4, 12). The severity of the workforce deficit makes changes, such as the increased level of clinical autonomy, associated with independent prescribing an attractive option to commissioners who seek to address gaps in service delivery. As the world leader in extending prescribing rights to nurses, pharmacists and allied health professions the findings are of significant importance to international policy makers who seek to learn from the pioneering advancement of prescribing rights in UK (31, 34) to inform their own approach to addressing the workforce deficit.

Internationally it is now common for physiotherapists, nurse practitioners, pharmacists, social workers, and psychiatric nurses to be located within extended primary care teams (87) with plans to extend this further recently announced (12, 18). Nearly 50% of appointments in UK general practice are for example, already provided by non-medical staff, i.e. nurses, pharmacists and allied health professionals (12, 15). Although in the current study, there were only four unplanned re-consultations for the same problem across the whole sample, these all related to patients (3 from NPs, 1 from IP) seeking further pain relief. While it is important to recognise that such small numbers do not provide an accurate basis for drawing conclusions about differences between IP and NP, it could be argued that the number of NP patients having to go back for pain medication was more than that for IPs and if this figure was multiplied across the total number of NP consultations per year it would actually be very costly in terms of additional GP appointments. This is important as the current deficit in primary care looks set to continue (88), with a recent proposal for home visits to be removed from the GP contract, and a government pledge to create 50 million more GP appointments year by 2024/25 (88, 89). As the third largest workforce in health and care in England, AHPs have great potential to contribute to transforming care, and ensuring ongoing access to medicines in these challenging times (18).

Non-medical prescribers see on average 5-30 patients per day (62, 90); based on a median of 17.5 consultations per day recent estimates indicate therefore that the current population of 90,000 qualified NMPs could potentially provide patients access to medicines in 1.56 million consultations per day, 580.35 million per year (90). Economic evaluation undertaken elsewhere in the UK indicates NMPs issue on average 2 medicines per day, thus facilitating the availability of 180,000 medicines via NMP per day, 65.7 million per year (90). However, it is important to note for the benefits of initiatives, such as prescribing, to be fully realised it is crucial that that they are supported by skilful implementation (12, 91, 92).

The economic evaluation of PP-IP is particularly important, given that identifying a sustainable solution that improves the worldwide deficit of health workers and makes best use of limited resources is imperative in ensuring ongoing access to medicines (12, 18). Our cost appraisal from the case sites suggest that PP-IP care delivery is more resource intensive than NP-PP. This arises through longer consultation duration, more ordering of medicines and tests (PO) and more discussions with colleagues (PTs). These costs, however, need to be considered in relation to benefits, many of which could not be measured in this study. Only a limited economic analysis was possible meaning that the findings should be treated with caution. Whilst the original intention had been to undertake a patient level micro costing analysis, data deficiencies limited what could be included. Further research is required to understand how team configurations affect care delivery, patient outcomes and costs.

The most complete data were available for consultation duration, and the calculation of associated costs showed IPs to incur slightly higher consultation costs than NP in both the PO and PT groups (£8.62 and £7.95 respectively). It is important to note however that consultation duration and associated costs may simply be driven by professional differences and clinic practices. The complexity of these arrangements means that the differences in cost could equally reflect service differences which would exist regardless of IP status. For example, the time spent in discussion with colleagues may reflect the multi-professional service that many case sites provided. Multi-professional, or team-working is a fundamental component of health care delivery in the UK and central to current government policy (93–95). There is increasing emphasis on establishing systems, rather than single episodes of care, that dissolve traditional boundaries (96, 97) to support the increasing number of people with long-term conditions.

There is limited evidence available with which to compare our study findings (32, 41, 51, 52, 91). Despite positive findings that NMP is safe, and provides beneficial clinical outcomes (32, 34, 79), the impact on the health economy, as reported in two recent systematic reviews examining clinical and cost effectiveness, remains unclear (51, 52, 91). The authors, as in this study, highlight the difficulty in separating NMP effects from the contributions of healthcare team members, and a lack of adequately powered randomised controlled trials examining NMP across clinical specialities, professions and settings (31, 51). Given that extended prescribing rights to nurses, pharmacists and AHPs offers a sustainable approach to improving the global workforce deficit, there is an urgent need to establish economic benefits, or otherwise of non-medical prescribing to inform future international policy developments. A different approach, involving highly targeted specific outcomes, and or longitudinal studies is therefore required. The development of a minimum data set of important outcome measures for NMP assessment would as Noblet et al. suggests (51), be highly beneficial, and generate the required evidence to evaluate the overall benefit of NMP and inform future developments in the UK and around the world.

### Strengths and limitations

In the first study to explore AHP prescribing, the 14 case sites supported an in-depth evaluation and comparison of PP-IP to PP-NPs in a range of care settings. Use of multiple methods of data collection, including an observational component, strengthens the trustworthiness of the findings. PP-IP participants were selected from a larger sample (n=35) who completed a trainee PP-IP survey and indicated that they would be willing to be involved in further research (62).

Despite challenges in matching sites, given the diversity of service settings, roles, and patient needs, between and within the two professions, patient characteristics indicated good matching on most factors. However, there are limitations and methodological challenges associated with using the same evaluation measures on two different professional groups for whom separate measures might have been more appropriate. The economic analysis was constrained as described above. An analysis of effectiveness was not possible because it was not feasible to collect data on specific indicators for change across the wide variety of conditions treated within PP consultations. Our ability to link each of the various aspect of patient data (i.e. observation, questionnaires, record audit) was also very limited as patients had the option to select which aspects of data collection they agreed to. As a result, it was not possible to match patients across the different data sets, or to complete some of the intended analysis.

## Conclusions

This study provides new knowledge about physiotherapist and podiatrist independent prescribing, the high level of care and patient satisfaction they provide. Given that extending prescribing responsibilities to nurses, pharmacists, and allied health professionals is increasingly being recognised as effective way to alleviate shortfalls in the global health workforce and ensure ongoing access to prescription medicines around the world this is important. PP-IP care delivery was found to be more resource intensive than NP-PP. However, this study is limited, and findings needs to be verified through further research, including a full economic analysis. A more focussed longitudinal exploration within each profession with targeted outcome measures would enable a more robust comparison of the impact of PP-IP across the United Kingdom, and inform further developments around the world.

## Acknowledgements

We would like to thank the physiotherapists and podiatrists, team members, patients and stakeholder participants who took part in our study, and our project advisory group and patient public involvement group for their advice and contributions throughout. Special thanks to assistance from Clareece Kerby, with data preparation and analysis, and Emma Konstantara with recruitment in private healthcare settings.

